# Distribution and characterization of *Aegilops cylindrica* species from Iran

**DOI:** 10.1101/525964

**Authors:** Behnam Bakhshi, Mohammad Jaffar Aghaei, Eissa Zarifi, Mohammad Reza Bihamta, Ehsan Mohseni Fard

## Abstract

Jointed goatgrass (*Aegilops cylindrica* Host; 2n = 4x = 28, C^c^C^c^D^c^D^c^) is a tetraploid remote relative of bread wheat (*Triticum aestivum* L; 2n=6x=42, AABBDD) with 2 genomes and 28 chromosomes. The diversity center of this species is in the Fertile Crescent and in central Asia and could also be found in many places in Iran. In this experiment, 359 accessions provided by National Plant Gene Bank of Iran (NPGBI) were used. Based on the geographical distribution, the highest distribution of *Ae. cylindrica* are from North, West and North West regions of Iran. The distribution data of *Ae. cylindrica* showed that the distribution centers in Iran are more than those reported in previous studies in Iran. Chromosome counting showed that all *Ae. cylindrica* accessions are tetraploid (2n=4x=28). Results of factor analysis for 9 morphological chromosome traits showed that karyotypic variation within accessions are related to the length of chromosomes and there is difference between accessions for their total chromosome length, but the karyotype of different accessions were almost the same for the symmetry. Low coefficient of variation in morphological traits as well as symmetric karyotypes of *Ae. cylindrica* species observed in this study could lead us to more confidently say that *Ae. cylindrica* could be a recently evolved species among remote relatives of bread wheat.

## 1. Introduction

Jointed goatgrass, originated from two species, is native to the Mediterranean, Middle East, Asia, and was also introduced to the Great Plains and the Pacific northwest of the United States (1, 2). It is a winter annual grass weed that infests over 3 million ha of winter wheat in the Pacific Northwest and Great Plains regions of the USA (3). It reduces winter wheat yields by interference and lowers harvested grain quality. Average yield loss with moderate to dense *Ae. cylindrica* infestations has been estimated to be 25% (4, 5) and it also has been estimated that the economic cost of *Ae. cylindrica* to winter wheat producers in the western United States is $145 million annually (6). Jointed goatgrass and winter wheat are closely related. Therefore, the development of selective herbicides to control this weed in winter wheat has been problematic.

*Aegilops cylindrica* is a bush type plant with 20-40 centimeters. It is characterized by narrow leaves (4-5 centimeters), none or very little hair, a narrow lanceolate spike (8 to 19 centimeters), almost ended with two incomplete spikelet. Each spike consists of 6-11 spikelets and breaks off entirely or disintegrates into segments. Each spikelet holds one to three seeds that are reddish-brown in color and reach maturity in mid-summer which is when the spikelets are fragile (1, 2, 4, 7) (Figure 1).

**Figure 1.**
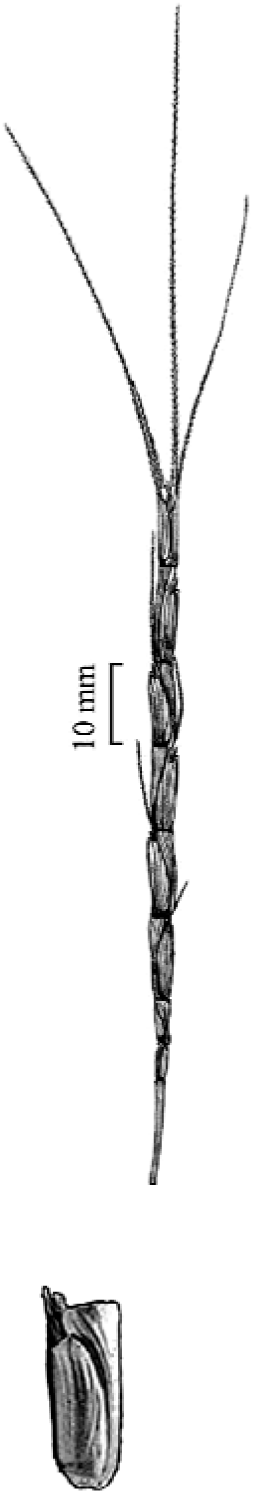
Spike and spikelet of *Ae. cylindrica* species.

The genomic constitution of *Ae. cylindrica* was determined by the analyses of chromosome pairing, storage proteins, isozymes, and differences in restriction length patterns of repeated nucleotide sequences (8). These studies identified the diploid species *Ae. caudata* L. (2n=2x=14, CC) and *Ae. tauschii* Coss. (2n=2x=14, DD) as the donor of the C and the D genome of *Ae. cylindrica*, respectively (8). Previously, the cytoplasm of *Ae. cylindrica* was shown to be contributed by *Ae. tauschii* (9, 10) but, more recent analysis with chloroplast microsatellite markers has shown that both *Ae. tauschii* (D-type cytoplasm) and *Ae. markgrafii* (C-type cytoplasm) have contributed their cytoplasms to *Ae. cylindrica* (11). High genome homology shared between *Ae. cylindrica* and it’s progenitor species and low intra-species polymorphism in *Ae. cylindrica* indicated it as a new species with little chromosome changes. The D genome chromosomes of *Ae. cylindrica* species are more similar to *Ae. tauschii* biotypes and D^cr2^ genome of hexaploid cytotype of *Ae. crassa* species than D genome of bread wheat (12, 13). These results indicate that there are different versions of D genomes for both *Ae. Cylindrica* and *T. aestivum* species.

This species has wide distribution from Western Europe to East Asia and even North America. The diversity center of this species is in the Fertile Crescent and in central Asia and could also be found in many places in Iran (1). *Ae. cylindrica* has spread westward to Greece, Bulgaria, Romania, Kosovo, Montenegro, Serbia, and Hungary. To the east, *Ae. cylindrica* is found in central Asia. Northwards, it is present in the Caucasus region and along the Black Sea coast. Though rare, this species is also present in the western arc of the Fertile Crescent involving Lebanon, Jordan, Syria, northern Iraq, and northwestern Iran. *Ae. cylindrica* is also adventives in many parts of Europe, Asia, and America (1).

The geographic distribution of *Ae. cylindrica* encompasses and extends beyond areas, where its diploid progenitors, *Ae. tauschii* and *Ae. markgrafii*, can be found (Figure 2).

**Figure 2.**
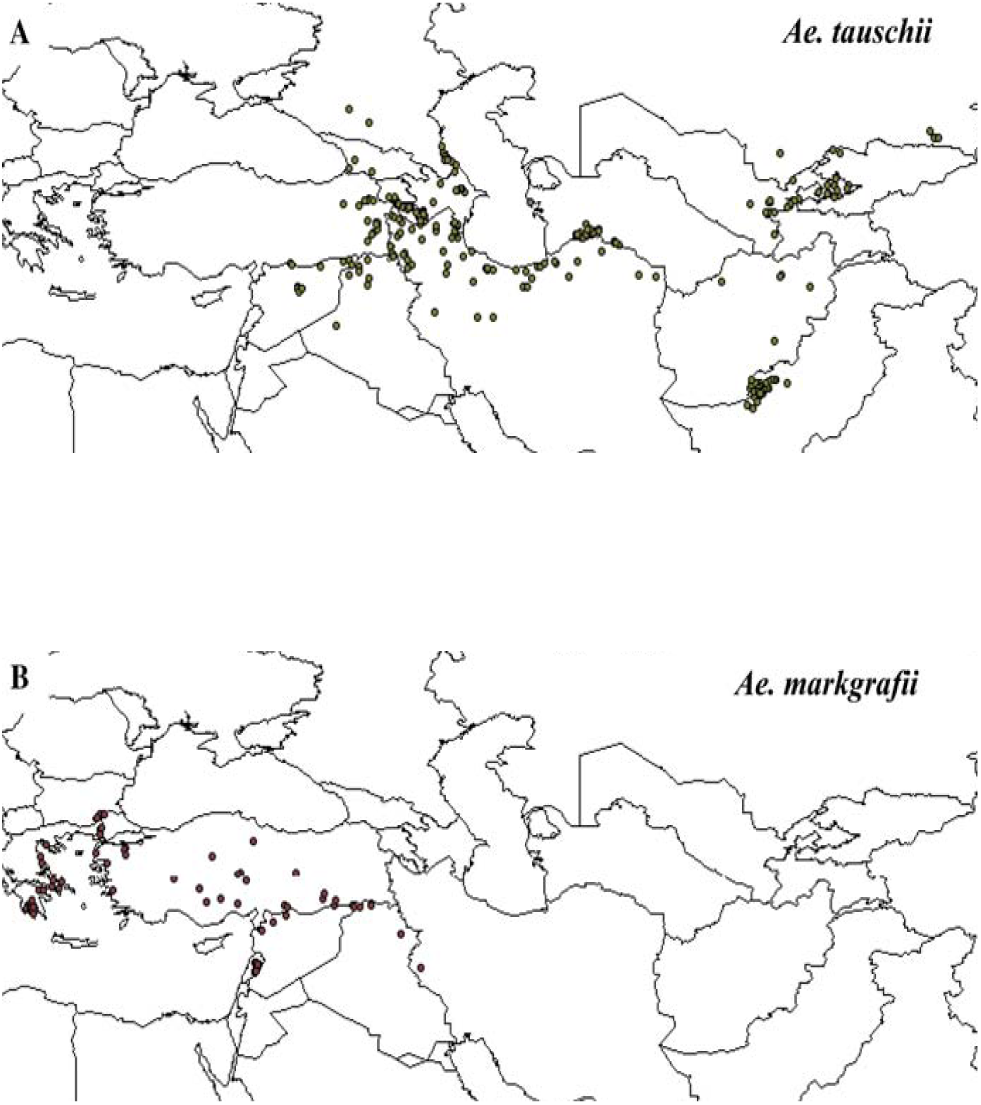
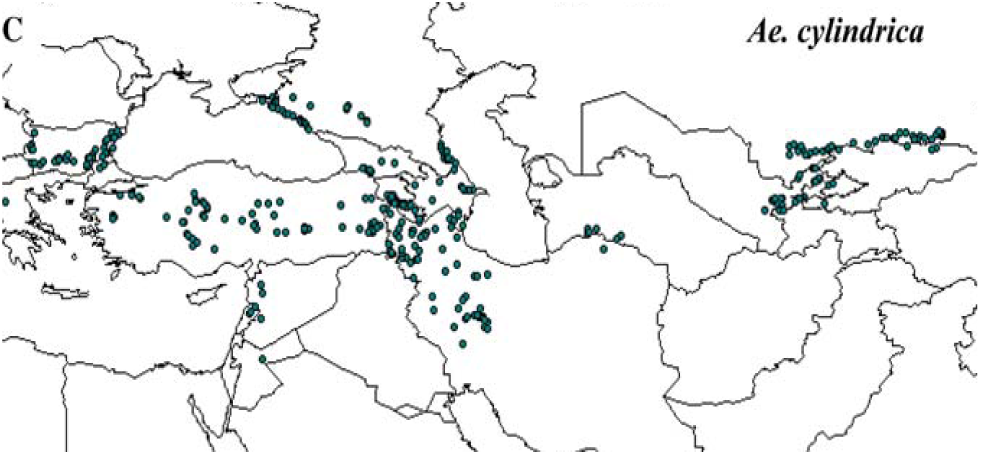
Maps showing the distribution of collections of *Ae. tauschii* (a), *Ae. markgrafii* (b), and *Ae. cylindrica* (c). The geographic coordinates were obtained from the system-wide information network for genetic resources (SINGER; http://singer.cgiar.org/).

Throughout its range of distribution, *Ae. cylindrica* is considered a weedy species, particularly in common wheat fields, where it chronically infests fields in the Mediterranean, the Middle East, Europe, and the United States of America (USA) (1, 3, 14). Jointed goatgrass has also been suggested as a source of genetic variation for wheat improvement (15-17) because it is a close relative of common wheat—both species carry the D genome donated by *Ae. tauschii* (18, 19). In addition, natural hybridization between wheat and jointed goatgrass suggests a potential for gene flow between these species under field conditions (20, 21). Thus, there is considerable interest in understanding various aspects of the evolution of *Ae. cylindrica* for its better management and use. Cytogenetic and molecular-based analyses suggested that the D genomes of *Ae. cylindrica* and *T. aestivum* were contributed by different biotypes of *Ae. tauschii* (12, 13). The D genome of hexaploid wheat has been shown to be more closely related to the D genome of *Ae. tauschii* ssp. *strangulata* than to *Ae. tauschii* ssp. *Tauschii* (22-24), whereas the D-type plastome and the D genome of *Ae. cylindrica* are more closely related to *Ae. tauschii* ssp. *tauschii* than to *Ae. tauschii* ssp. *strangulate* (11).

Although molecular genetic diversity and ploidy level of *Ae. tauschii* has been reported before (25, 26), no extensive study has been done to identify cytogenetic and morphologic characteristics of this species. We have collected many accessions from different regions of Iran that it could be remarkable to identify potential genetic diversity among these accessions. In this study, we analyzed chromosomes features of *Ae. cylindrica* as well as distribution and morphological characteristics of this species because of wide distribution of this species in Iran. Since this species is a relative of bread wheat it is important to identify available genetic diversity in this species to be used in necessary condition in bread wheat breeding programs.

## 2. Materials and Methods

### 2.1. Plant material

359 accessions were used in this experiment, which were provided from National Plant Gene Bank of Iran (NPGBI). These accessions were collected from sixteen provinces of Iran (West Azarbaijan, East Azarbaijan, Ardebil, Zanjan, Qazvin, Kurdistan, Hamedan, Kermanshah, Ilam, Lorestan, Chaharmohal Bakhtiari, Mazandaran, Tehran, Esfahan, Semnan and Khorasan). Traits and measuring methods are shown in the Table 1. Evaluation of all traits was conducted using three replications of each accession. The mean and mode was calculated for quantitative and qualitative traits, respectively. Estimated statistical parameters traits were calculated for quantitative traits. Shannon and Weaver diversity index were calculated for measuring qualitative traits. Non-standard values of diversity index (Hc) and standard diversity index (SDI) was calculated as follows(27):

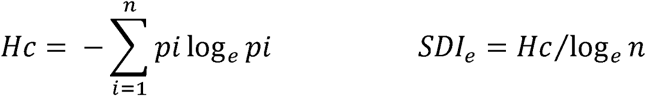

**Table 1.** Traits and methods of measure.

| Rows | Traits | Measuring methods |
| --- | --- | --- |
| 1. | Plant height | Measuring stem length per plant in cm |
| 2. | Maturity date | Counting the number of days from 21 March to 50% of plant maturity |
| 3. | Flowering date | Counting the number of days from 21 March to 50% of plant Flowering |
| 4. | Spikelet seed number | Counting the number of seed per spikelets (per plant) |
| 5. | Leaf number of rachis | Counting the number of stem leaves (per plant) |
| 6. | Spikelet number per spike | Counting the number of spikelets (per spike) |
| 7. | Node number of rachis | Counting the number of stem nodes (per plant) |
| 8. | Kernel length | Length measurement of seeds (per plant) (mm) |
| 9. | Spike length | Measuring spike length (per plant) (cm) |
| 10. | Spikelet length | Spikelet length measurement (per plant) (mm) |
| 11. | Length of spikelet glume | Measuring length of spikelet glume (per plant) (mm) |
| 12. | Kernel width | Measuring kernel width (per plant) (mm) |
| 13. | Spikelet width | Measuring spikelet width (per plant) (mm) |
| 14. | Width of spikelet glume | Measuring spikelet glume width (per plant) (mm) |
| 15. | Rachis width | Measuring rachis width (per plant) (mm) |
| 16. | Spike width | Measuring spike width (per plant) (mm) |
| 17. | Seed tissue | Named grain flour, half glass and glass, with the numbers 3, 5 and 7, respectively |
| 18. | Stamen color | Named white, yellow and brown stamens, with numbers 1, 2 and 3, respectively |
| 19. | Seed color | Named white, red and purple grains, with numbers 1, 2 and 3, respectively |
| 20. | Glume color | Named white, red to brown and purple to black with numbers 1, 2 and 3, respectively |
| 21. | Spike-axis fragility | Named very fragile, moderate and strong spike with numbers 1, 2 and 3, respectively |
| 22. | Growth habit | Named standing, half standing and asleep plants with numbers 1, 3 and 5, respectively |
| 23. | Glume hairs | Named glabrous, with less cracking, medium and high Glume hairs with numbers 1, 3, 5 and 7, respectively |

In this formula, for certain traits, such as c, n, including the number of phenotypic classes and Pi is equal to the frequency of bushes.

### 2.2. Chromosome counting

Based on the technique, developed at International Maize and Wheat Improvement Center (CIMMYT) institute (28), root tips were collected between 9 AM to 10:30 AM, and then placed in a Petri dish, on a filter paper moistened with α-bromonaphthalene pre-treatment solution. The samples were pre-treated about 2.5 to 3.5 hours, but generally 3 hours as pre-treatment time which was used in this study, resulting satisfying chromosome contraction and high mitotic index. After pre-treatment, the root tips were transferred to vials, containing 0.2% aceto-orcein and refrigerated (4°C), until they are being used. Afterwards, the root tips were transferred to 2% aceto-orcein, in order to intensify the staining for 2 days before squashing. After staining the aceto-orcein was removed from the vial and 45% acetic acid was added to fill about a quarter of the vial. Vial was heated over a flame to bring the contents to a slow boiling. After boiling, the vial contents (45% acetic acid + root tip) were transferred into an evaporating dish. A root tip was taken from it and placed over on filter paper to remove extra acetic acid. Apical root tip measuring 2-2.5mm was cut and placed on dry microscope slide. The root tip was squashed by an arrow-head needle, and a small drop of 45% acetic acid was quickly added to the squashed tissue. The slide was then slightly warmed and a cover glass was placed gently over on the macerated cellular area. The cover glass slides were gently dabbed with coarse filter paper, the slide was heated slightly, placed between folded filter paper on a flat surface and thumb pressure applied directly to the cover glass. After squashing, the slide was suitable for observing chromosomes by microscope.

### 2.3. Karyotype preparation

In order to prepare karyotypes of *Ae. cylindrica*, 12 accessions of *Ae. cylindrica* were used (Table 2).

**Table 2.** Accessions of *Ae. cylindrica* used in karyotype study.

| Province | City | Accession No. | Longitude |  | Latitude |  |
| --- | --- | --- | --- | --- | --- | --- |
|  |  |  | Degree | Seconds | Degree | Seconds |
| West Azarbaijan | Naghadeh | 50 | 45 | 22 | 36 | 57 |
| Lorestan | Borujerd | 96 | 57 | 20 | 37 | 28 |
| Zanjan | Zanjan | 363 | 48 | 29 | 36 | 40 |
| Kermanshah | Songhor | 286 | 47 | 34 | 36 | 47 |
| East Azerbaijan | Maragheh | 406 | 46 | 16 | 37 | 24 |
| East Azerbaijan | Hashtrud | 408 | 47 | 4 | 37 | 28 |
| Ilam | Shirvan | 312 | 46 | 34 | 33 | 46 |
| Ardabil | Ardabil | 332 | 48 | 17 | 38 | 15 |
| Kermanshah | West Islamabad | 379 | 46 | 32 | 34 | 7 |
| East Azerbaijan | Urmia | 45 | 45 | 2 | 37 | 32 |
| Kermanshah | Javanrud | 381 | 46 | 22 | 35 | 3 |
| Hamadan | Hamadan | 393 | 48 | 31 | 34 | 48 |

### 2.4. Study of Karyotypes

Chromosomes are named according to the location of centromere in the chromosome (26). Comparison between karyotypes in different accessions of a species was performed by comparing their symmetry in this study. In addition, the Stebbins method for determining the degree of symmetry has been used. Stebbins (1971) believes this as suitable method to recognize the role of karyotypes in evolution of the species. In addition, total form percentage index (TF) (29), the relative percentage of the longest chromosome to the shortest chromosomes (S) (30), coefficient chromosomal length variations (CV) (31), the average ratio of long to short arm (R) (30) and range of chromosome length variation (V) (32) were calculated in this study. Furthermore, evaluation of karyotype evolution was calculated using Dispersion Index (DI) to show a few differences that were not visible in Stebbins indicators (33). Factor analysis for morphological features of chromosomes based on principal component analysis and Varimax rotation was also conducted in this study. For measuring different parts of the chromosomes and for analyzing morphological data of chromosomes, Micromeasure software (34) and the SPSS software were used, respectively.

## 3. Results and Discussion

### 3.1. Geographical distribution of of *Ae. cylindrica* accessions in Iran

Investigation on collecting location and geographical distribution of *Ae. cylindrica* accessions reveales that this species predominantly grows in the range of 800 to 2000 meters altitude. Thus this species is adapted to mountainous ecosystems and not to low altitude ecosystems of Caspian shores, southern shore, Khuzestan and Ilam. Also, the results of the geographical distribution using ILWIS software also showed that the highest distribution of *Ae. cylindrica* was from that North, West and North West regions, including East Azarbaijan, West Azarbaijan and Kermanshah provinces contrasting to the southern and south-east regions that showed the lowest distribution (Figure 3). However, it may be found anywhere on the mountainous areas of Alborz and Zagros. Some populations could even be found on briny margins of Uromieh lake and Semnan, Northern Khorasan and around Qom. Whereas, *Ae. cylindrica* doesn’t grow on salty deserts of central and southern Iran. While, it is abundant in central Iranian land, *Ae. cylindrica* is usually found in the more elevated northern strip of the central Iranian desserts and not in the more arid region of southern and central areas.

**Figure 3.**
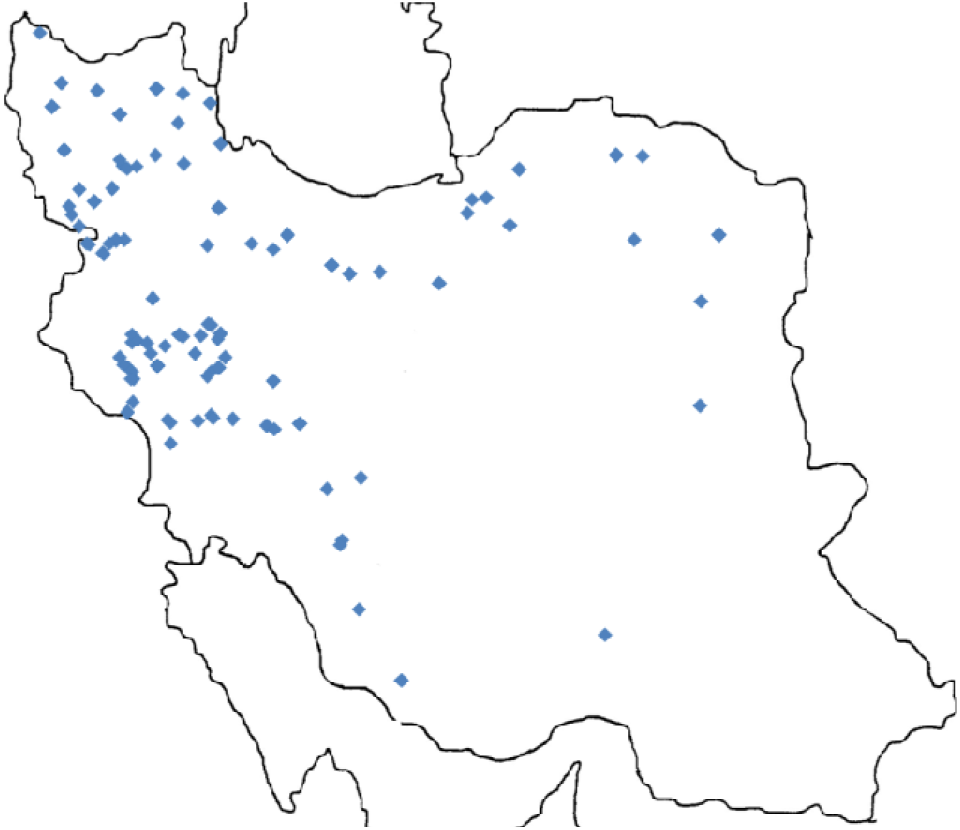
Geographical distribution of *Ae. cylindrica* in Iran.

The present investigation on the geographical distribution of *Ae. cylindrica* showed that the distribution centers are more than reported in a previous study (11) in Iran. In the present study, in addition to north, west and northwest, northeast and southwest of Iran have also been identified as distribution centers for this species. The results also showed that this species mostly present in mountain ecosystems and it is not found at low altitude ecosystems such as the margins and the southern coast ecosystems of Caspian Sea. Previous studies indicated that the evolution of bread wheat occurred in high altitudes of Caspian Sea ecosystems (35, 36). On the other hand the results of this study showed that the *Ae. cylindrica* species has a wide distribution in this region. With this explanation, the possibility of challenging unproven hypothesis that *Ae. cylindrica* had a potential to be as a donor of hexaploid wheat D genome (37) might be more acceptable.

### 3.2. Statistical parameters in the studied traits of *Ae. cylindrica* species

Rachis width, spikelet seed number, plant height, kernel width and leaf number of rachis traits showed the highest phenotypic coefficient of variation, with 13.14, 11.33, 10.85, 10.43 and 10.11 percent, respectively. Most of these traits have been also observed among traits with high diversity in other collected accession in Iran including *Aegilops tauschi* and *Aegilops crassa* (38, 39). Furthermore, maturity date, spikelet length and length of spikelet glume traits indicated the lowest phenotypic coefficient of variation with 4.80, 5.02, and 5.61 percent. Most standard deviations were also related to plant height, flowering date and maturity date and the lowest standard deviation were related to rachis width, spikelet width and width of spikelet glume (Table 3). White and flour form kernel, brown and glabrous glumes, moderate fragility spikes, brown stamens and standing bushes were predominantly observed traits in the research field. Furthermore, growth habit showed the most variation using non-standard and standard diversity index of Shannon-Weaver (40). Thus, according to Shannon-Weaver index, growth habit could be introduced as the most effective qualitative trait to distinguish *Ae. cylindrica* populations (Table 4). Relatively low level of phenotypic variation coefficients for different traits were obtained in this study showing that this species is relatively new. However, it has been observed high genetic diversity for *Aegilops tauschi* in northern area of Iran (38).

**Table 3.** Statistical parameters for quantitative traits evaluated in the collection of *Ae. cylindrical*.

|  | Mean | Mod | Middle | Minimum | Maximum | Range of Variation | Standard deviation | Variance | Coefficient of variation |
| --- | --- | --- | --- | --- | --- | --- | --- | --- | --- |
| Plant height | 59.67 | 59.00 | 60.33 | 38.33 | 89.67 | 51.33 | 6.47 | 41.88 | 10.85 |
| Flowering date | 65.12 | 67.00 | 66.00 | 58.00 | 93.00 | 35.00 | 4.34 | 18.82 | 6.66 |
| Maturity date | 89.23 | 87.00 | 88.00 | 58.00 | 98.00 | 40.00 | 4.28 | 18.32 | 4.80 |
| Leaf number of rachis | 3.46 | 3.33 | 3.33 | 2.67 | 4.33 | 1.67 | 0.35 | 0.12 | 10.11 |
| Spikelet seed number | 2.04 | 2.00 | 2.00 | 1.33 | 3.00 | 1.67 | 0.23 | 0.05 | 11.33 |
| Spikelet number per spike | 9.28 | 9.00 | 9.33 | 6.67 | 11.67 | 5.00 | 0.85 | 0.73 | 9.18 |
| Node number of rachis | 2.92 | 3.00 | 3.00 | 2.00 | 3.67 | 1.67 | 0.21 | 0.04 | 7.14 |
| Kernel width | 7.08 | 6.83 | 7.07 | 5.47 | 9.10 | 3.63 | 0.52 | 0.27 | 7.39 |
| Spike length | 8.65 | 9.00 | 8.67 | 6.00 | 10.33 | 4.33 | 0.74 | 0.54 | 8.53 |
| Spikelet length | 11.72 | 11.73 | 11.73 | 8.20 | 14.47 | 6.27 | 0.59 | 0.35 | 5.02 |
| Length of spikelet glume | 9.28 | 10.17 | 9.83 | 8.23 | 11.23 | 3.00 | 0.55 | 0.30 | 5.61 |
| Kernel width | 2.33 | 2.40 | 2.33 | 1.63 | 5.63 | 4.00 | 0.24 | 0.06 | 10.43 |
| Spikelet width | 2.69 | 2.67 | 2.67 | 2.27 | 3.63 | 1.37 | 0.17 | 0.03 | 6.51 |
| Width of spikelet glume | 2.88 | 2.87 | 2.87 | 2.40 | 4.27 | 1.87 | 0.19 | 0.04 | 6.70 |
| Rachis width | 1.27 | 1.17 | 1.27 | 0.83 | 1.90 | 1.07 | 0.17 | 0.03 | 13.14 |
| Spike width | 2.82 | 2.83 | 2.80 | 2.30 | 4.30 | 2.00 | 0.22 | 0.05 | 7.67 |

**Table 4.**
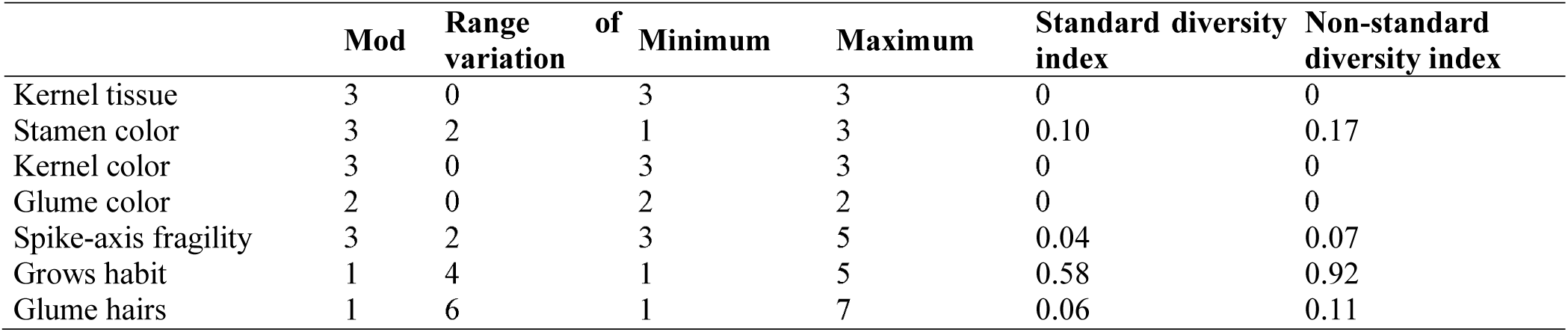
Statistical parameters for qualitative traits evaluated in the collection of *Ae. cylindrical*.

### 3.3. Karyotype analysis of accessions

Chromosome counting showed that most of *Ae. cylindrica* accessions are tetraploid (2n = 4x = 28). Furthermore, cytogenetic studies showed no aneuploids and B chromosomes, but difference in chromosome length is shown in this study. Collecting place and accession No are presented in Table 2 and chromosomes count, satellites count in karyotype and also karyotypic formula are shown in Table 5. The metaphase cell and ideogram of various accessions are presented in figures 4 to 15. All investigated accessions had a satellite in the short arm of chromosomes No. 8. The presence of satellite in the same pair of chromosomes is also reported (41). Considerable variation was observed in total length of chromosomes (TLC) and the average length of chromosomes (C). High variation in length of chromosomes may be a sign of genome adaptation of this species to those places from where they have been collected. The largest chromosome (11.33 micrometer) was found in Zanjan and the smallest chromosome (4.87 micrometer) was observed in the accession of Urmia and both of them were sub-metacentric. Maximum and minimum of long arm to short arm ratio was observed in Islam Abad Gharb (2.129) and in Shirvan (1.78) accessions, respectively. For total form percentage index (TF), the maximum (35.32) and minimum (31.18) TF was observed in Ardabil and Zanjan accessions, respectively. This data shows that karyotypes of Ardebil and Zanjan accessions have the highest and the lowest symmetry. The highest coefficient of variation (CV) was found in accession from Naghadeh and the lowest coefficient of variation was found in the Javanrud accession. All of the accessions that were collected from the north-west and the West of Iran, belonged to A2 position of the Stebbins Table, indicating a relatively symmetrical karyotype for recently evolved species with short evolutionary history. Also, distribution index of chromosome (DI) showed that the Hashtrud accession had the highest DI (7.43) and Zanjan accession had the lowest DI (18.5). This observation indicated that the Hashtrud accession had the highest symmetry in contrast to Zanjan accession. DI could be more reliable than other indicators because three important karyotypic criteria, including; variation in chromosomes length, centromere position and relative size of chromosomes are involved in DI calculation (Table 6).

**Table 5.** Collecting place, chromosomes count, satellites count and karyotypic formula of *Ae. cylindrical*.

| Collecting place | Chromosomes count | Satellites count | Karyotypic formula |
| --- | --- | --- | --- |
| Naghadeh | 28 | 1 | 2m* + 6sm* + 6st* |
| Borujerd | 28 | 1 | 1m + 8sm + 5st |
| Zanjan | 28 | 1 | 2m + 6sm + 6st |
| Songhor | 28 | 1 | 4m + 4sm + 6st |
| Maragheh | 28 | 1 | 2m + 9sm + 3st |
| Hashtrud | 28 | 1 | 1m + 9sm + 4st |
| Shirvan | 28 | 1 | 5m + 8sm + 1st |
| Ardabil | 28 | 1 | 3m + 9sm + 2st |
| West Islamabad | 28 | 1 | 3m + 6sm + 5st |
| Urmia | 28 | 1 | 1m + 12sm + 1st |
| Javanrud | 28 | 1 | 1m + 10sm + 3st |
| Hamadan | 28 | 1 | 2m + 7sm + 5st |

\* m; Metacentric chromosome, sm; Submetacentric chromosome and st; Subtelocentric chromosome

**Table 6.** Karyological characteristics of *Ae. cylindrica* accessions.

| Collecting place | Stebins Index | Distribution index of chromosome | CV of chromosome length | Centromere index | Minimum of short arm to long arm ratio percentage | Total length of chromosomes (TLC) | Average of short arm to long arm ratio | Average of chromosome length | Total length of chromosome | Range of chromosome length variation |
| --- | --- | --- | --- | --- | --- | --- | --- | --- | --- | --- |
| Naghadeh | 2A | 6.46 | 13.48 | 0.32 | 65.01 | 32.36 | 2.09 | 8.68 | 121.61 | 3.77 |
| Borujerd | 2A | 5.84 | 12.09 | 0.32 | 66.51 | 32.54 | 2.07 | 7.57 | 105.97 | 3.12 |
| Zanjan | 2A | 5.19 | 11.45 | 0.31 | 68.42 | 31.19 | 2.20 | 9.22 | 129.16 | 3.57 |
| Songhor | 2A | 5.65 | 11.90 | 0.32 | 67.77 | 32.21 | 2.10 | 7.46 | 104.46 | 2.95 |
| Maragheh | 2A | 6.41 | 12.84 | 0.33 | 67.13 | 33.31 | 2.00 | 8.43 | 118.12 | 3.41 |
| Hashtrud | 2A | 7.44 | 15.06 | 0.33 | 60.76 | 33.07 | 2.02 | 7.70 | 107.88 | 3.87 |
| Shirvan | 2A | 7.25 | 12.88 | 0.23 | 65.38 | 32.19 | 1.78 | 9.13 | 127.89 | 3.89 |
| Ardabil | 2A | 6.12 | 11.19 | 0.35 | 69.74 | 35.33 | 1.83 | 6.76 | 94.64 | 2.49 |
| West Islamabad | 2A | 6.10 | 12.99 | 0.31 | 64.19 | 31.96 | 2.13 | 8.27 | 115.87 | 3.71 |
| Urmia | 2A | 5.93 | 11.15 | 0.34 | 69.07 | 34.73 | 1.88 | 5.85 | 81.94 | 2.18 |
| Javanrud | 2A | 5.62 | 11.04 | 0.33 | 69.75 | 33.75 | 1.96 | 6.76 | 94.68 | 2.43 |
| Hamadan | 2A | 6.61 | 12.98 | 0.33 | 65.82 | 33.71 | 1.97 | 6.45 | 90.38 | 2.77 |

**Figure 4.**
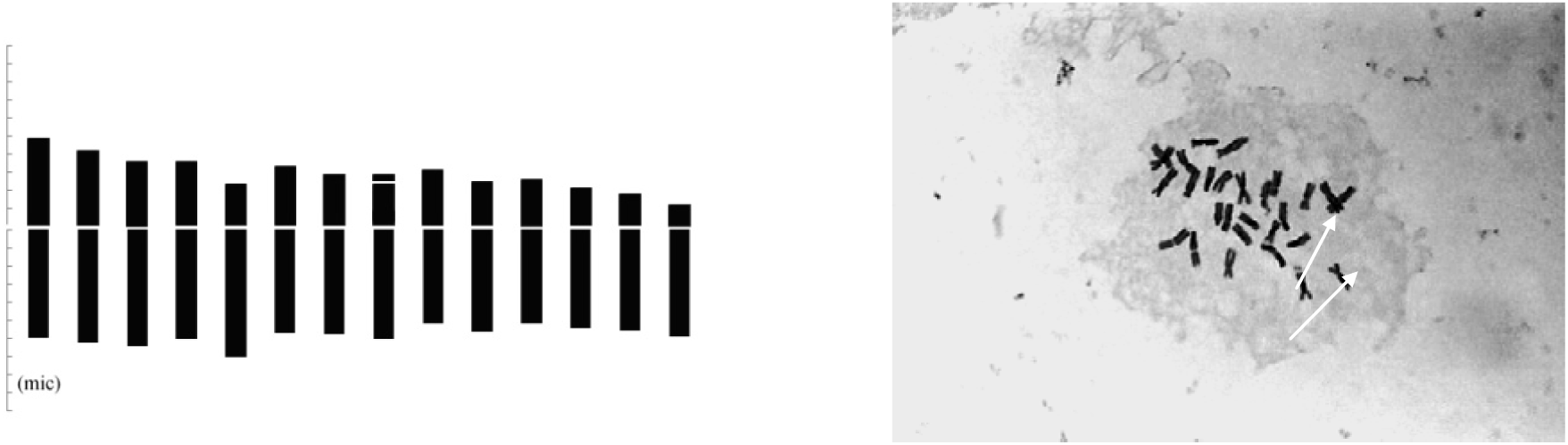
metaphase chromosomes picture and ideogram of Naghadeh population

**Figure 5.**
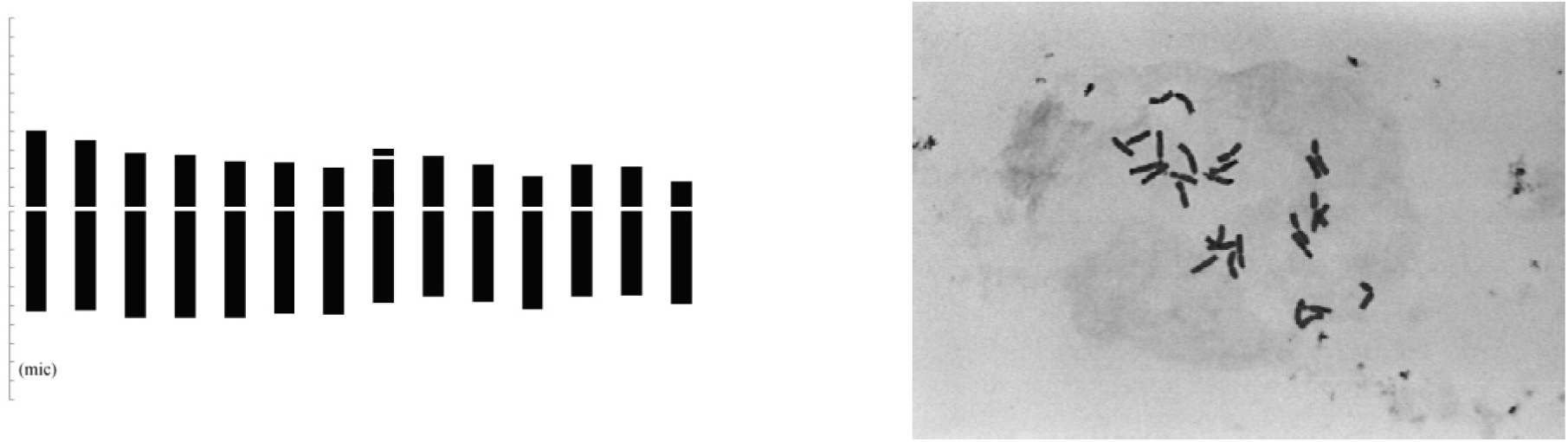
metaphase chromosomes picture and ideogram of Broujerd population

**Figure 6.**
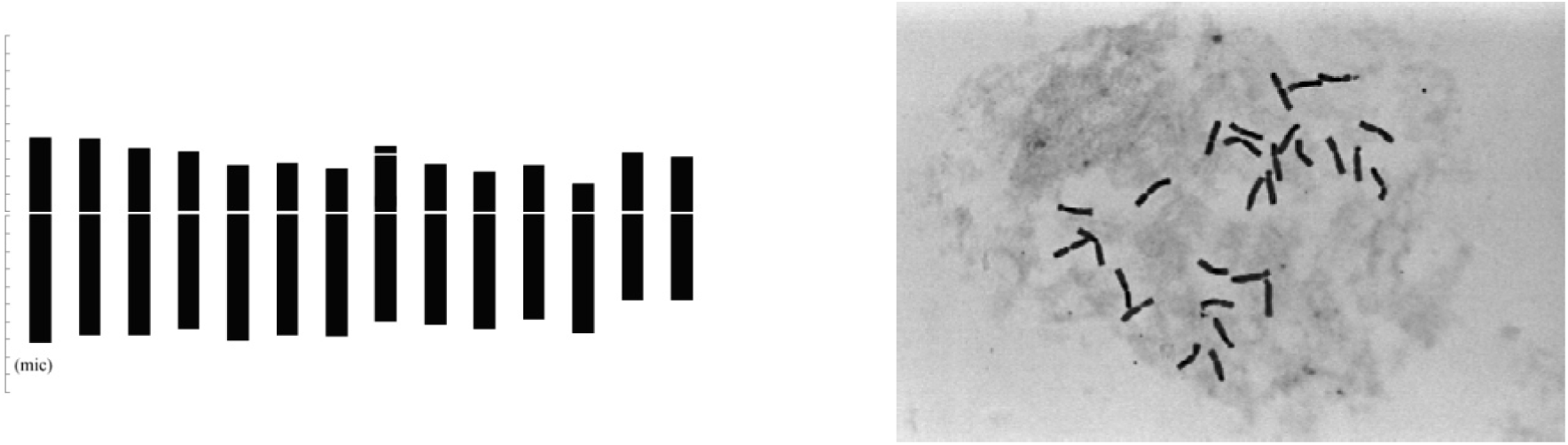
metaphase chromosomes picture and ideogram of Zanjan population

**Figure 7.**
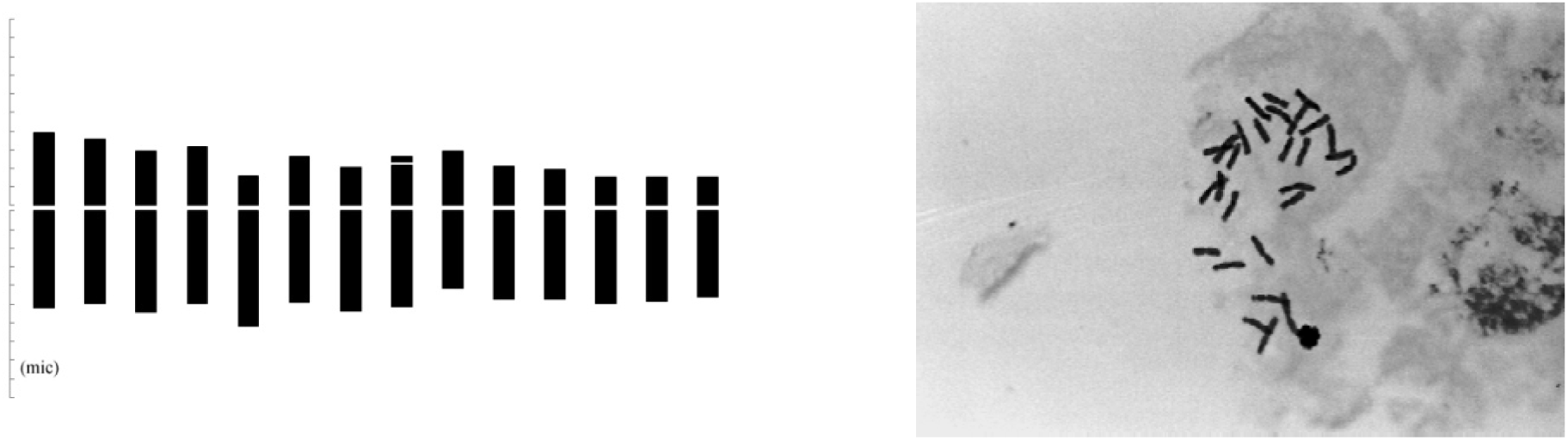
metaphase chromosomes picture and ideogram of Songhor population

**Figure 8.**
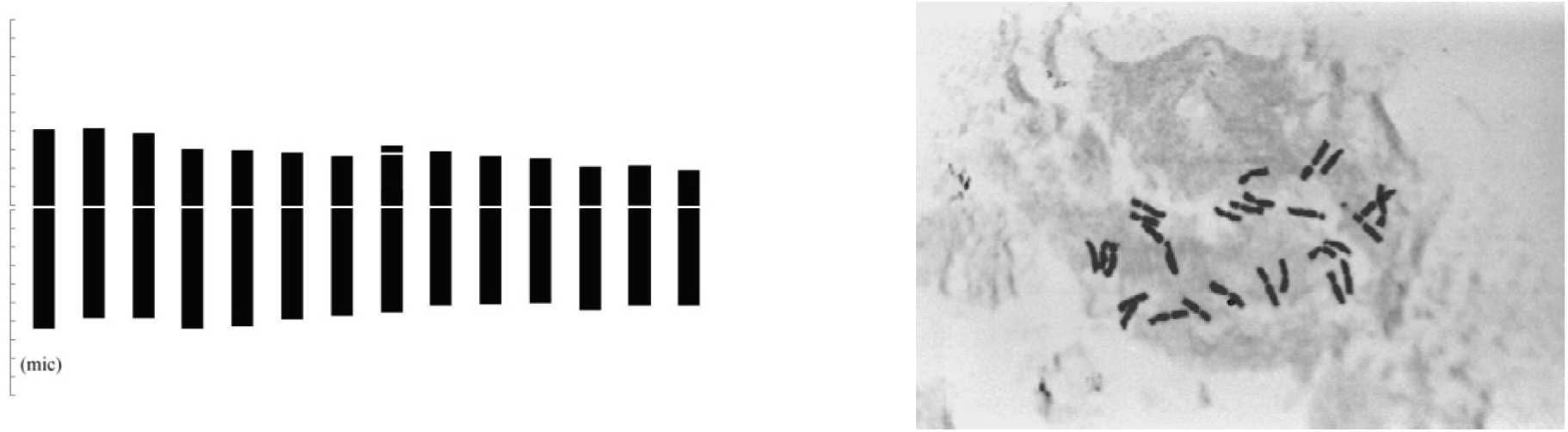
metaphase chromosomes picture and ideogram of Maragheh population

**Figure 9.**
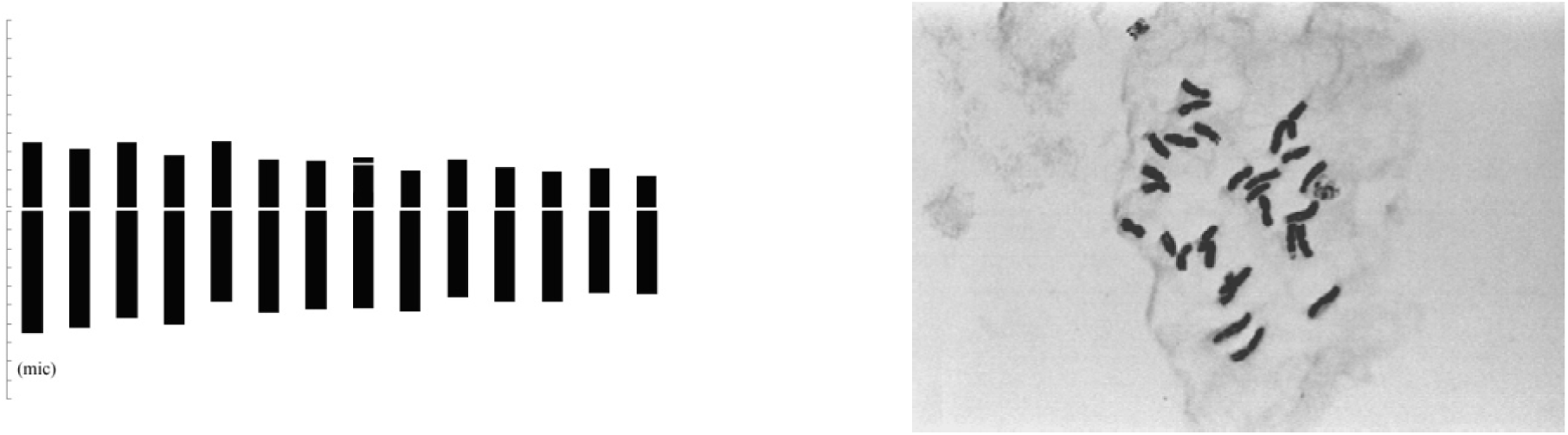
metaphase chromosomes picture and ideogram of Hashtroud population

**Figure 10.**
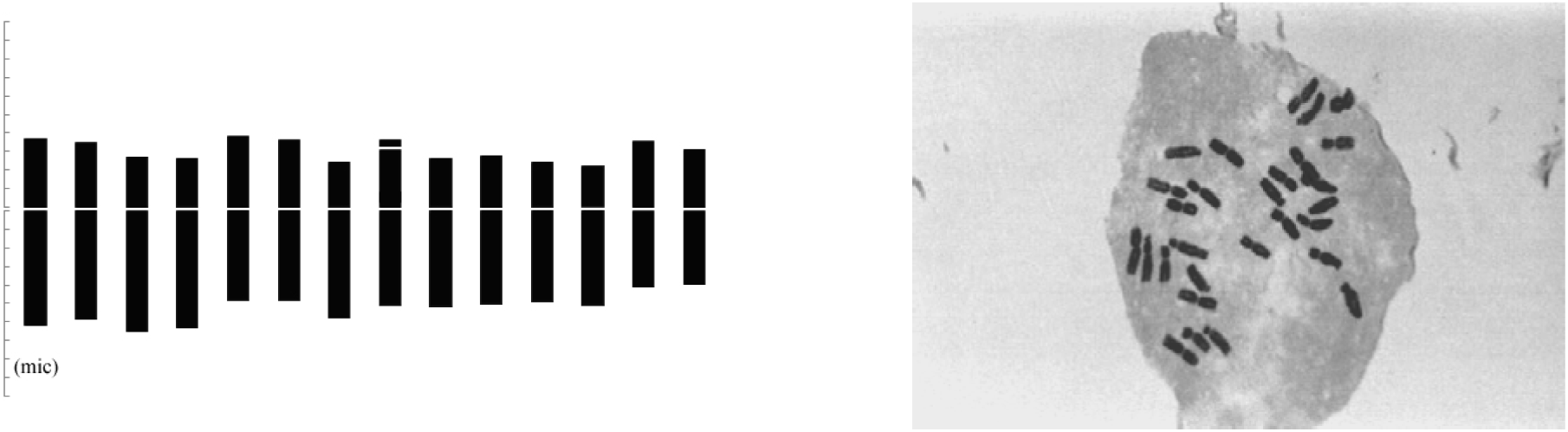
metaphase chromosomes picture and ideogram of Shirvan population

**Figure 11.**
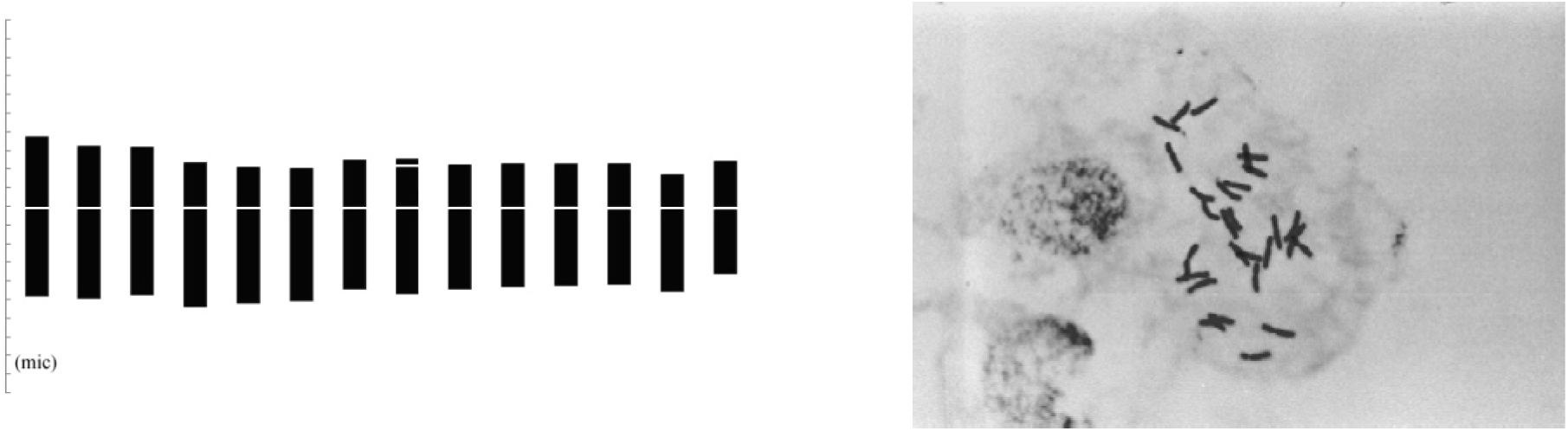
metaphase chromosomes picture and ideogram of Ardebil population

**Figure 12.**
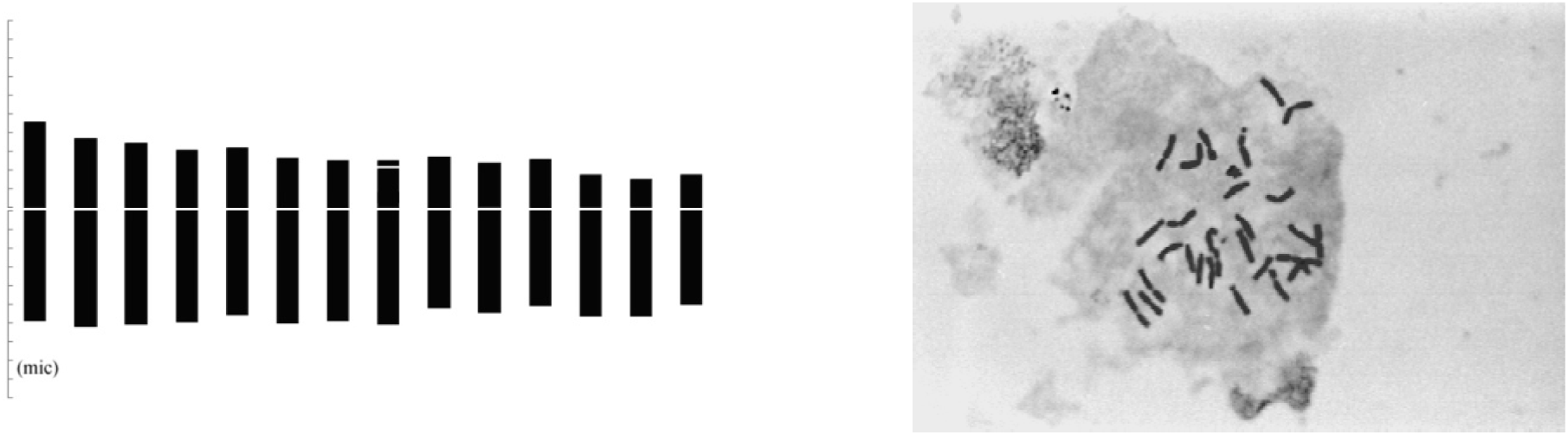
metaphase chromosomes picture and ideogram of Islam Abad Gharb population

**Figure 13.**
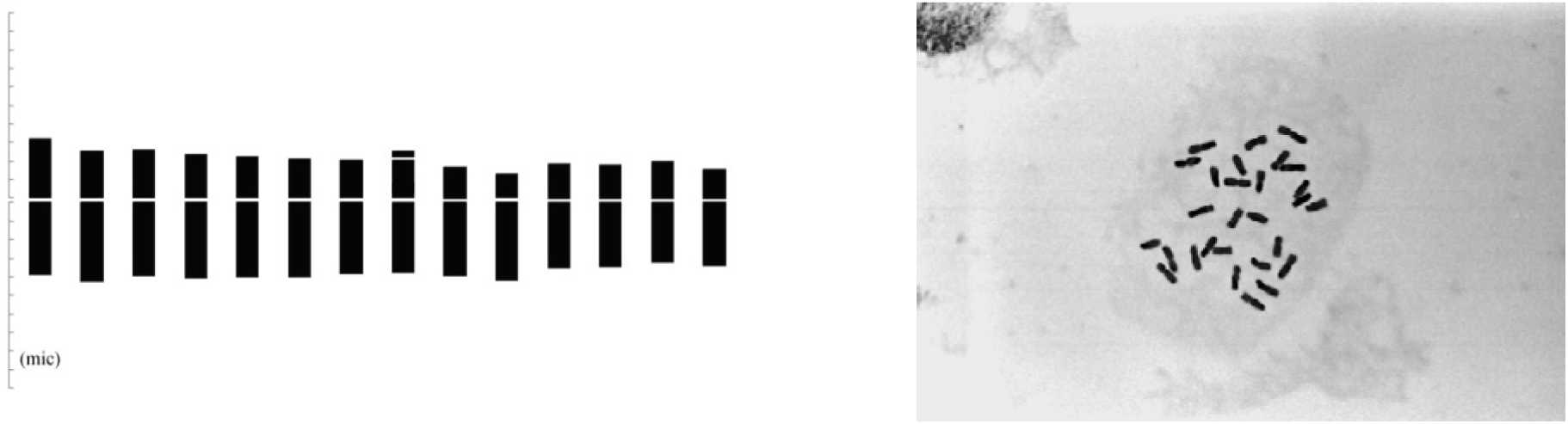
metaphase chromosomes picture and ideogram of Orumia population

**Figure 14.**
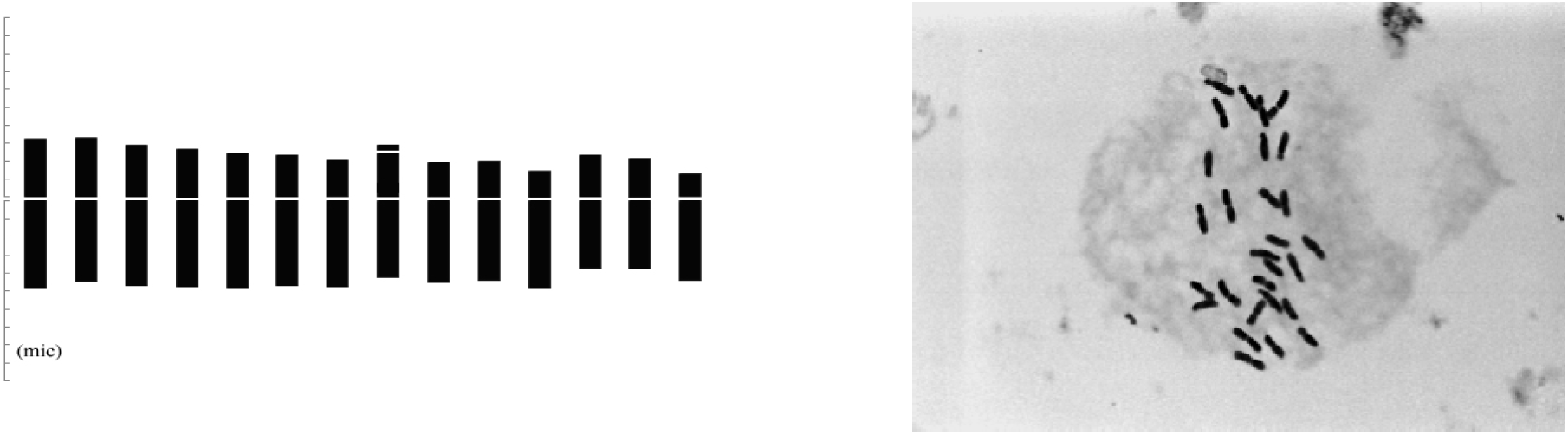
metaphase chromosomes picture and ideogram of Javanroud population

**Figure 15.**
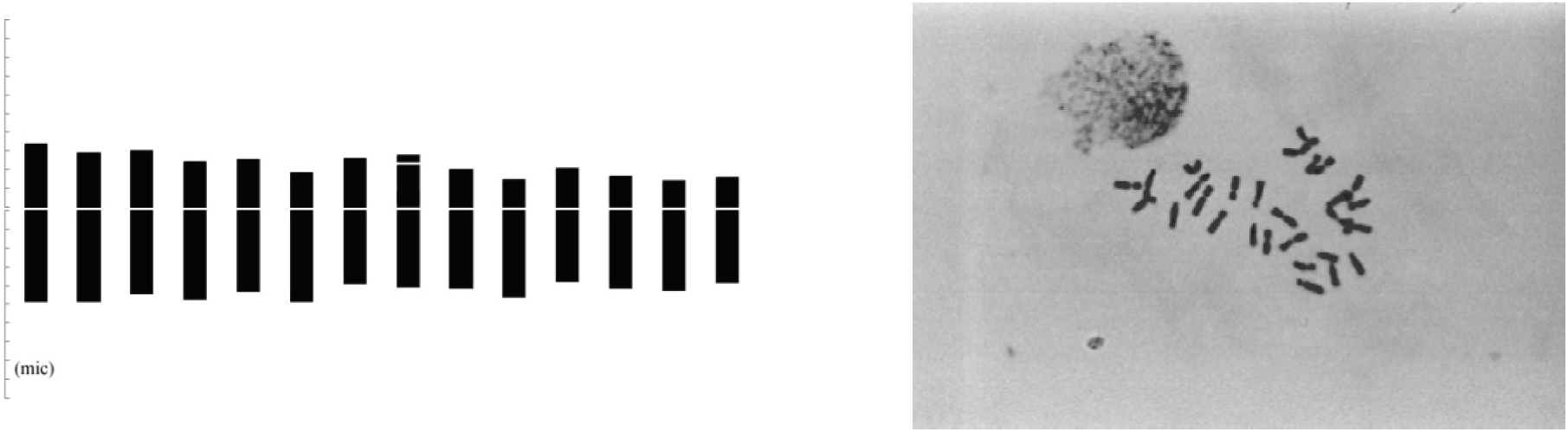
metaphase chromosomes picture and ideogram of Hamedan population

The results of factor analysis of 9 chromosomal morphology traits showed that the first two factors had Eigen values greater than one with 65.83 to 88.39 percent variance and these two factors were responsible for diversity in the accessions (Tables 7 & 8).

**Table 7.** Eigen values, percentage of variance and cumulative variance factor.

| Factors | Eigen values | Percentage of variance | Cumulative variance factor |
| --- | --- | --- | --- |
| 1 | 5.93 | 65.83 | 65.83 |
| 2 | 2.03 | 22.56 | 88.39 |

**Table 8.**
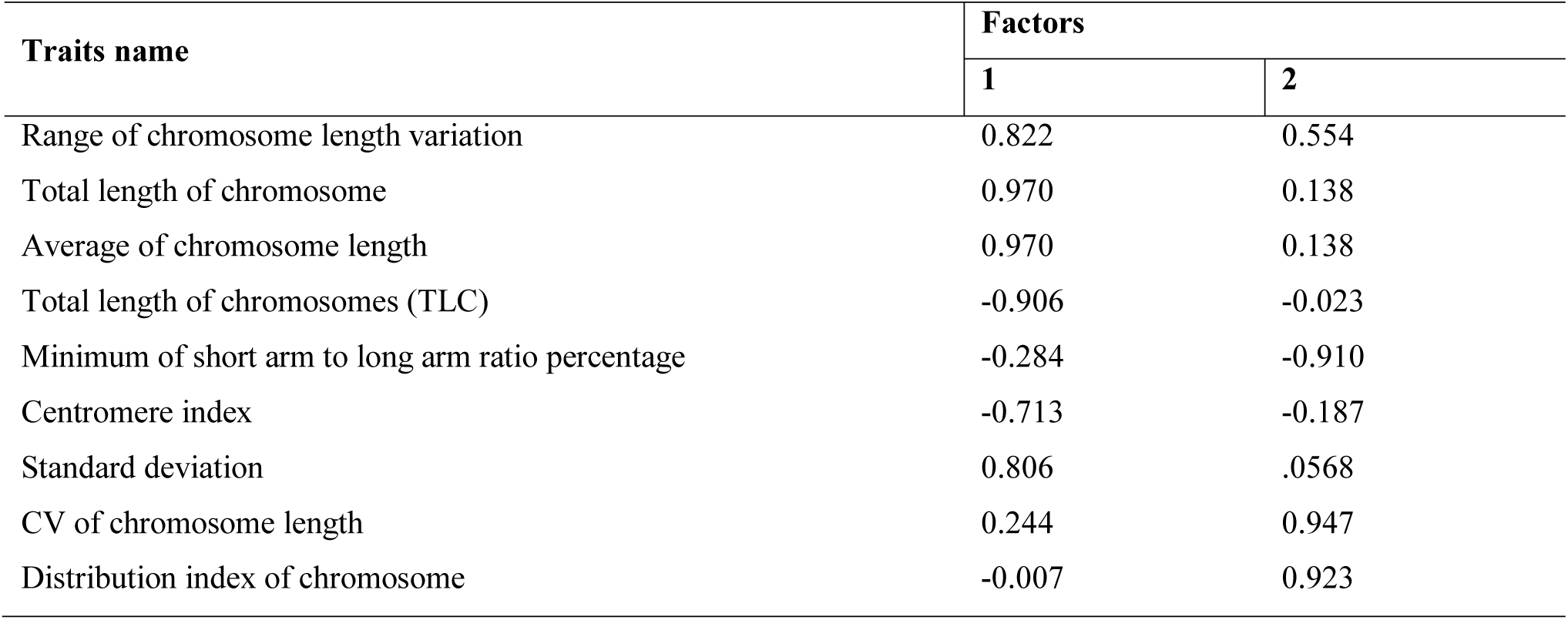
The first two factors derived from factor analysis for morphological traits of chromosomes.

| Traits name | Factors |  |
| --- | --- | --- |
|  | 1 | 2 |
| Range of chromosome length variation | 0.822 | 0.554 |
| Total length of chromosome | 0.970 | 0.138 |
| Average of chromosome length | 0.970 | 0.138 |
| Total length of chromosomes (TLC) | -0.906 | -0.023 |
| Minimum of short arm to long arm ratio percentage | -0.284 | -0.910 |
| Centromere index | -0.713 | -0.187 |
| Standard deviation | 0.806 | .0568 |
| CV of chromosome length | 0.244 | 0.947 |
| Distribution index of chromosome | -0.007 | 0.923 |

Range of chromosome length variation, total length of chromosome, average of chromosome length, total length of chromosomes (TLC), centromere index, and standard deviation of chromosome length traits had the largest factor coefficients in the first factor. Because most of these traits depend on the chromosomes length, this factor is named length of chromosomes. This factor presents 65.83 percent of the total variance that shows there is great diversity for traits related to chromosome length among accessions. In the second factor, CV of chromosome length, minimum of short arm to long arm ratio percentage and distribution index traits had the largest factor coefficients. Thus, the second factor was named as karyotype symmetry duo to all of these traits indicated karyotype symmetry of the accessions. The second factor accounted for 22.55 percent of the total variance, indicating that there was no great difference between accessions in terms of symmetry and confirming the placement of accessions at the 2A position of the Stebbins table. Therefore, the results of factor analysis show that karyotypic variation within accessions is related to the length of chromosomes and there is difference between accessions for their total chromosome length; but the karyotype of different accessions are same for their symmetry and they are relatively symmetrical. As morphological studies were conducted, low coefficient of variation coupled with symmetric karyotype indicates *Ae. cylindrica* as a recently evolved species.

## 4. Conclusion

*Aegilops cylindrica* diversity centers are mostly located in the Northwest regions where the highest numbers of collection sites are distributed. We also observed that *Ae. cylindrica* accessions of Iran should be treated as a recently evolved species due to low diversity in morphological traits and symmetric karyotypes.

